# Estimating wildlife vaccination coverage using genetic methods

**DOI:** 10.1101/129064

**Authors:** Freya Smith, Andrew Robertson, Graham C. Smith, Peter Gill, Robbie A. McDonald, Gavin Wilson, Richard J. Delahay

## Abstract

Vaccination is a potentially useful approach for the control of disease in wildlife populations. The effectiveness of vaccination is contingent in part on obtaining adequate vaccine coverage at the population level. However, measuring vaccine coverage in wild animal populations is challenging and so there is a need to develop robust approaches to estimate coverage and so contribute to understanding the likely efficacy of vaccination.

We used a modified capture mark recapture technique to estimate vaccine coverage in a wild population of European badgers (*Meles meles*) vaccinated by live-trapping and injecting with Bacillus Calmette-Guérin as part of a bovine tuberculosis control initiative in Wales, United Kingdom. Our approach used genetic matching of vaccinated animals to a sample of the wider population to estimate the percentage of badgers that had been vaccinated. Individual-specific genetic profiles were obtained using microsatellite genotyping of hair samples which were collected both directly from trapped and vaccinated badgers and non-invasively from the wider population using hair traps deployed at badger burrows.

We estimated the percentage of badgers vaccinated in a single year and applied this to a simple model to estimate cumulative vaccine coverage over a four year period, corresponding to the total duration of the vaccination campaign.

In the year of study, we estimated that between 44-65% (95% confidence interval, mean 55%) of the badger population received a vaccine dose. Using the model, we estimated that 70-85% of the total population would have received at least one vaccine dose over the course of the four year vaccination campaign.

This study represents the first application of this novel approach for measuring vaccine coverage in wildlife. This is also the first attempt at quantifying the level of vaccine coverage achieved by trapping and injecting badgers. The results therefore have specific application to bovine tuberculosis control policy, and the approach is of significance to the wider field of wildlife vaccination.

## Introduction

Vaccination can contribute to disease control by reducing the number of susceptible and/or infectious individuals in a population and hence reduce the number of new infections. The approach has been applied to the control of wildlife disease reservoirs (Blancou *et al.* 2009), particularly oral vaccination of red foxes (*Vulpes vulpes*), which resulted in the eradication of rabies from much of Central and Western Europe by the end of the 20th century (Müller *et al.* 2015). More recently, oral vaccination of wild boar has made a significant contribution to the control of Classical Swine Fever in parts of Europe (Rossi *et al.* 2015).

The success of vaccination as a method of disease control is largely determined by two factors, (1) vaccine efficacy in an individual and (2) vaccine coverage across a population. Attempts to estimate vaccine coverage in wild animals typically rely on post-vaccination surveillance in order to detect direct or indirect markers of vaccination in the target species (e.g. Rosatte *et al.* 2008, Rossi *et al.* 2015). This approach is usually dependent on the capture or recapture of vaccinated animals (demanding significant investments of time and resources), or the collection of carcasses (e.g. from hunting (Johnston *et al.* 1988)). Consequently, it can be difficult to estimate coverage, particularly for species that are not hunted. Consequently, there is a clear demand for novel, effective methods for estimating vaccine coverage in wildlife populations.

Bovine tuberculosis (bTB) is a chronic disease of cattle caused by *Mycobacterium bovis* (*M. bovis*). In many European countries, control measures focused on cattle, principally test and slaughter based on the tuberculin skin test, have been successful at reducing and/or eliminating infection from cattle. However, in places where infection persists within a wildlife reservoir, disease control is more complex. This is the case in the UK and Ireland, where the European badger (*Meles meles*) is the principal wildlife reservoir for bTB and a potential source of infection for cattle (Krebs *et al.* 1997, Bourne *et al.* 2007, Godfray *et al.* 2013).

Bacillus Calmette–Guérin (BCG) is an attenuated strain of *M. bovis* that has been used as a live vaccine in badgers. When delivered by intramuscular injection BCG has been shown to slow the progression of disease, reducing both the severity of lesions and the excretion of bacilli in experimentally challenged captive animals (Chambers *et al.* 2011, Lesellier *et al.* 2011). Field trials have also shown a significant reduction in positive serological test results in wild badgers vaccinated with BCG (Chambers *et al.* 2011, Carter *et al.* 2012), consistent with reduced infection and/or disease progression. Finally, Carter *et al.* (2012) have demonstrated an indirect beneficial effect of vaccination, whereby unvaccinated cubs born into badger social groups with a higher proportion of vaccinated adults were significantly less likely to test positive for *M. bovis* infection. Together, these studies provide evidence that sustained vaccination could bring about a reduction in the prevalence of *M. bovis* in badgers (Chambers *et al.* 2014). Furthermore, mathematical modelling suggests that this could have a positive impact on levels of infection in cattle (Smith *et al.* 2012).

Although oral vaccination may offer the most practical strategy for widespread deployment of BCG in badgers in the long term (Blancou *et al.* 2009), an oral BCG bait has not yet been licensed and is unlikely to be available for a number of years (Chambers *et al.* 2014). On the other hand, an injectable BCG vaccine, BadgerBCG, was licensed by the UK Veterinary Medicines Directorate in 2010 and is currently available for the purpose of vaccinating trapped badgers by intramuscular injection (Brown *et al.* 2013). Field deployment of injectable BadgerBCG has now been carried out at a number of locations across England and Wales (APHA 2015). Badgers are live-captured in mesh traps, injected with BCG and temporarily marked by hair clipping and the application of coloured stock marker. Given the field logistics of vaccination in a disease control setting, as opposed to a research trial, there are currently no estimates of the level of vaccine coverage that has been achieved during these badger vaccination initiatives. Furthermore, there are no existing estimates of badger trapping efficiency that are directly applicable to vaccination. For example, Smith and Cheeseman (2007) estimated that trapping efficiency in ‘proactive’ cull areas during the Randomised Badger varied between 35-85%. However, the pattern and intensity of badger trapping carried out during the RBCT has not been reproduced by vaccination programmes during which traps can be set for a maximum of two consecutive nights as a condition of licences for trapping badgers for vaccination in England and Wales.

We estimated vaccine coverage during one year of a badger vaccination campaign in Wales, UK, where badger vaccination had been introduced as part of measures aimed at reducing the local incidence of bTB in cattle. We used a genetic mark recapture approach (trap sample matching), to match vaccinated animals to a genetically ‘marked’ sample from the wider population on the basis of badger hair genotype profiles. We then combined our results with estimates of population turnover in order to calculate the percentage of the total badger population that could be expected to have received at least one vaccine dose after four years of vaccination (the total duration of the vaccination campaign).

## Methods

### Study area and population

The bTB Intensive Action Area (IAA) is a 288 km^2^ area of high bTB incidence in cattle, primarily located in north Pembrokeshire, Wales. Since 2010, the IAA has been subject to additional disease control measures over and above those in place in the rest of Wales, with the aim of reducing and eventually eliminating bTB in cattle in the area. The suite of measures applied to the IAA includes badger vaccination, by live-trapping and injection, which was initiated in 2012 (Welsh Government 2012) and has subsequently been repeated on an annual basis. In each of the first three years of the programme, over 1300 doses of BCG were administered (in excess of 4000 doses in total) to individual badgers from a population of unknown size (Welsh Government 2016). This vaccination programme was entering its fourth and final year at the time the present study took place.

### Badger trapping and vaccination

For the purposes of vaccination, badgers were captured using steel mesh traps which were placed close to active badger setts or on active runs and baited with peanuts. Access for trapping was authorised by land owners for 249 km^2^ or 86% of the IAA (Welsh Government 2016). Where access to setts or runs was not permitted, traps were located near linear features and boundaries (e.g. hedges) of accessible land. Traps were set for two, typically consecutive nights (unless interrupted by adverse weather conditions) following a period of approximately 7 days during which traps were pre-baited but not set. Captured badgers were vaccinated by intra-muscular injection of 1 ml of prepared BadgerBCG (BCG Danish strain 1331 vaccine, Statens Serum Institut, Copenhagen, Denmark) administered through the mesh of the trap. In order to avoid dosing animals more than once in the same year, vaccinated animals were temporarily marked by clipping the fur and applying a coloured stock marker prior to release. Vaccination was organised into seven cycles of three to four weeks duration, each targeting a different portion of the IAA and scheduled at approximately monthly intervals between May and October (Welsh Government 2016).

### Genetic sampling

For vaccinated animals, hair samples for identification were taken by plucking a sample of approximately 5-10 guard hairs from the rump of every trapped individual using artery forceps. Hairs were placed in a labelled sample bag together with a sachet of desiccant (Minipax ® absorbent packets, Sigma-Aldrich).

For the wider badger population, we deployed ‘hair traps’ at a sample of main setts following a methodology developed for estimating badger abundance (Frantz *et al.* 2004, Scheppers *et al.* 2007, Judge *et al.* 2017). Hair traps consist of strands of barbed wire suspended across the sett entrance holes or on nearby badger runs. As badgers pass under the traps, their guard hairs catch on the wire barbs and can be collected for analysis.

As in previous studies, main setts were used as a proxy for badger social groups because there is usually only one main sett per badger social group territory (Cresswell *et al.* 1990, Wilson *et al.* 1997). Hair traps were deployed (n=560) at a sample of 72 (28%) main setts from a total of 260 main setts identified during previous badger surveys in the IAA. The number of hair traps deployed per sett ranged from 2-18 (median eight) depending on the number of entrance holes and badger runs found. Setts were selected by random sampling, stratified by scheduled vaccination month (‘trap round’) and revisited prior to setting hair traps. Setts that at this time were inactive, inaccessible or deemed not to be a main sett were substituted by randomly selected replacements from the same trap round.

Hair traps remained *in situ* for at least 4 weeks and were visited on alternate days over a 28 day sampling period (Scheppers *et al.* 2007). On each visit, all the hairs on a given trap were removed and collected into a labelled bag containing a sachet of desiccant. If hair had caught on multiple barbs of the same trap the hairs from each barb were placed in separate sample bags, and then placed together within a labelled bag representing the specific hair trap on that day of collection. Once samples had been collected, hair traps were decontaminated by brief exposure to a naked flame.

### Genetic typing

Hair samples were temporarily stored at room temperature (vaccinated animal samples, maximum 14 days) or in vehicles (hair trap samples during 28 day collection period followed by maximum 14 days at room temperature) prior to being refrigerated at 4°C. Up to ten hairs were selected from every labelled sample bag, each pool of hairs representing either an individual vaccinated animal or a specific hair trap-day combination. Hairs were selected on the basis of the size of the follicle as DNA recovery is expected to be more successful from larger follicles. In the case of the hair trap samples, all hairs selected originated from a single barb. Hairs from the remaining barbs were retained and were only analysed if the profile resulting from the original sample was suspected of being of mixed origin, see below.

DNA was extracted with a suspension of chelex resin (Frantz *et al.* 2004) using the Qiagen DNeasy® Blood and Tissue Kit. Genetic profiles were obtained by amplifying ten microsatellites (*Mel-103*, *Mel-104*, *Mel-105*, *Mel-107*, *Mel-110*, *Mel-113*, *Mel-114*, *Mel-115*, *Mel-116* and *Mel-117;* (Carpenter *et al.* 2003)). Microsatellite fragments were detected on an Applied Biosystems 3730xl Genetic Analyser and were analysed and sized using GeneMapper® Software (version 5). Incomplete profiles (where amplification had failed at one or more loci) were excluded from the analyses.

Allele calling (designation of genotypes) was performed automatically using the Applied Biosystems GeneMapper® software. Each genotype was then reviewed manually by two operators. DNA profiles generated from the hair trap samples underwent a further manual review in order to screen for DNA profiles which contained contributions from more than one animal as a consequence of pooling sampled hairs (mixed profiles). Suspected mixed profiles were identified on the basis of the presence of more than two alleles at one or more loci and/or a difference in peak height between heterozygous alleles such that a minimum threshold of heterozygote balance (the smallest allele in peak height divided by the largest allele in peak height) was exceeded at one or more loci (S1 Appendix). These profiles were excluded from further analyses in order to avoid artificially inflating the number of unique profiles obtained from the background population. Where possible, we repeated extraction and genotyping of these samples (this time, based on individual hairs rather than pooled hairs, if necessary, from a different barb to the original sample analysed), although this was not always possible because the entire hair sample had often been used up in the first round of analysis.

Genotype data were subsequently checked for the presence of null alleles (alleles which failed to amplify reliably for a particular microsatellite) using the programme CERVUS (Marshall *et al.* 1998). The output indicated that null alleles were present at microsatellite *Mel-116* (+0.2703). As a result, this microsatellite was not used for trap sample matching.

### Trap sample matching

Genotypes derived from captured and vaccinated badgers were matched to the those from the background population using the statistical package ALLELEMATCH (Galpern *et al.* 2012), executed in R 3.0.2 (R Development Core Team 2016). This allowed us to estimate the proportion of the population that had been trapped and vaccinated. Profiles were matched at the nine remaining microsatellites. Two profiles were identified as being from the same animal if they shared at least 17 of the 18 available alleles. Matching of profiles that differ by one, or a small number of alleles is an approach that is commonly used in wildlife genetics studies (e.g. Hettinga *et al.* 2012, Judge *et al.* 2017) where the quantity and quality of DNA may be low resulting in more frequent genotyping errors and hence a greater potential for mismatching replicate samples.

We then adjusted the raw percentage of matched profiles for false positive errors (matching of profiles from different animals), false negative errors (mismatching of profiles from the same animal) and a movement rate for all setts within dispersal distance of the boundary of the IAA (to account for the fact that some individuals identified by hair trapping may have moved out of the area, prior to cage traps being deployed). See S2 Appendix for further details of trap sample matching including calculation of error and movement rates.

### Modelling cumulative vaccination coverage

We used a simple quantitative model to estimate the cumulative percentage of badgers likely to have been vaccinated in the IAA by the end of a four year campaign. Model assumptions included a fixed rate of vaccination (our estimate of the percentage of badgers trapped and vaccinated in 2015), and a fixed rate of survival (based on published estimates). The result represents the estimated percentage of the standing badger population in the IAA, i.e. all badgers alive in the area at the end of the 2015 vaccination year that can be expected to have received at least one vaccine dose during their lifetime. See S3 Appendix for further details of the model and model parameters.

## Results

Plucked hairs were obtained from badgers at all 1118 trapping and vaccinating events, of these samples, 1065 (95%) were successfully genotyped. Initial exploratory analyses utilised data from all 10 microsatellites. We identified forty pairs of identical genotype profiles amongst the plucked hair samples. These matches were consistent with recapture and revaccination of the same individuals (S2 Appendix) and duplicate genotypes were therefore excluded from further analyses. This resulted in a final dataset of 1025 genotype profiles from vaccinated badgers.

In total, 682 hair trap-day samples were collected from 66 out of 72 (92%) badger main setts, of which 384 yielded a complete genetic profile. Of these, 224 profiles were identified as individuals. The remaining 160 were identified as potential mixtures. Reanalysis of single hairs from 78 trap-day samples yielded a further 44 complete genetic profiles. Therefore, the total number of useable profiles from the background population was 268 (39% of all trap-day samples). These profiles represented 141 unique individual badgers, from 53 main setts. The number of unique individuals identified per sett (and hence per social group) ranged from 1 to 10 (median 2), S4 Appendix.

Of the 141 genotype profiles from hair traps, 68 (48%) matched those from hair samples from trapped and vaccinated badgers (i.e. were identical for at least 17 of 18 available alleles). Accounting for error and movement rates (S2 Appendix), the adjusted estimate of the percentage of badgers trapped and vaccinated in the IAA in a single year, i.e. 2015, was 55% (95% confidence intervals 44-65%) of the total population.

Assuming consistent annual coverage at the observed level, we estimate that by the end of a four year vaccination campaign 70-85% of the total population would have received at least one vaccine dose.

## Discussion

Wildlife vaccination can be a useful strategy for limiting transmission of infections within and between species, either as an additional or alternative to other means of disease control (Blancou *et al.* 2009). In order to optimise vaccine deployment, both in terms of cost and effectiveness, it is important to understand the level of coverage achieved. In the present study we demonstrate a modified capture mark recapture method to estimate coverage for a vaccine delivered by trapping and injection.

The badger vaccination project in the Welsh Government’s Intensive Action Area (IAA) provided the context for the present study. The programme of work in the IAA represents the largest badger vaccination campaign undertaken to date with more than 1000 doses of injectable BCG administered to badgers annually over a four year period (2012 to 2015). The trapping protocol employed during the programme is the same as that recommended under current guidelines for badger vaccination and is therefore broadly comparable with other vaccination programmes in England and Wales. In the present study we matched genotypes from vaccinated badgers to those from a background population to estimate the level of coverage achieved in the IAA during one year of the vaccination campaign.

Our results suggest that in a single year of vaccination, 44-65% of badgers in the IAA were trapped and vaccinated. Assuming equivalent levels of coverage were achieved in preceding years, we estimate that by the end of four consecutive years of vaccination, 70-85% of the total badger population in the IAA would have received at least one dose of vaccine.

An important assumption underlying all capture mark recapture methodologies is that the ‘marked’ sample of animals is representative of the target or wider, background population. In this study, the marked sample was established using remote sampling of hairs and genotyping. This approach has an advantage over conventional capture mark recapture methods because it means that sampling of the background population is not dependent on trapping animals and so uses an independent method from the sampling of the vaccinated population. As a result, our wider, hair-trapped sample may well have included trap-shy animals that could otherwise have been missed. We therefore expect this sample to be more representative of the background population. In order to further satisfy the assumption of representative sampling, the setts included in this study were randomly selected, and sample size was high (we deployed hair traps at more than a quarter of all main setts in the IAA). We also collected samples over a relatively long time period (alternate days for a four week period (see Scheppers et *al*. 2007) in order to sample animals that were not permanently resident at the main sett. However, sampling bias in favour of animals that were more active at main setts and spent less time at outlier setts, cannot be ruled out. If such animals were also more likely to be trapped for vaccination this could have resulted in an overestimate of coverage. It is also possible that sampling was biased towards adult badgers, as cubs may have passed beneath hair traps without coming into contact with the barbed wire. However as sampling did not take place until June, it will have avoided the main period when very small post-emergent cubs would be expected to be present (Roper 2010).

Mistyping of genotypes due to genotyping error can occur by a variety of different mechanisms (reviewed by Pompanon *et al.* 2005). We mitigated the risk of false negative matches (failure to match samples from the same animal) by matching at 17 alleles rather than the full complement of 18 alleles. This resulted in a number of presumed false matches between different vaccinated animals. However, this source of error was quantified and incorporated into our final estimate of coverage and so should not have biased the results.

Genotyping of DNA from pooled hairs could have contributed to mismatching by creating ‘false’ genotypes within the background population, by combining those of multiple individuals. To minimise the chance of this occurring, all hair-trap profiles were screened for markers of mixed DNA prior to matching. Our protocol for screening was conservative (S1 Appendix), such that we almost certainly rejected some genotypes from single animals, reducing the overall sample size. However, as these individuals should be random in relation to the likelihood of being caught, this should not have biased our estimate of coverage.

As BCG provides incomplete immunity in badgers (Chambers *et al.* 2014), the percentage of badgers captured and vaccinated cannot be taken to indicate the level of protection achieved in the population, particularly since a proportion of the population will already have been infected and the effect of post-exposure vaccination, although unknown, is assumed to be zero. In addition, the duration of immunity achieved by BCG vaccination of badgers has yet to be determined: in neonatal calves a temporal reduction in immunity has been observed (Thom *et al.* 2012), while in humans efficacy may reduce but is still long lasting (Nguipdop-Djomo *et al.* 2016). Some individuals will have received multiple doses over the course of the IAA badger vaccination campaign and it is also not known what effect this might have on the extent or duration of immunity in vaccinated animals. The significant protective effect observed in unvaccinated cubs living in social groups with vaccinated adults (Carter *et al.* 2012), should also be taken into account in translating our estimate of coverage into a meaningful estimate of possible protection.

Although there are uncertainties, badger vaccination may still be an effective management approach. Estimates of R0, the number of secondary infections per individual in a naïve population, are low (Smith 2001) which implies that relatively low levels of vaccination can drive the disease extinct, given enough time.

Over recent years, there has been increasing support for vaccination of wildlife against bTB, in an effort to reduce the risk of onwards disease transmission to cattle (Blancou *et al.* 2009). As the primary wildlife reservoir for *M. bovis* in the UK and Ireland, and a species increasingly recognised as playing a role in the persistence of *M. bovis* infection in mainland Europe (Gortazar *et al.* 2012), the European badger has been a particular focus for bTB wildlife vaccination research. After almost 30 years of research, captive and field studies have demonstrated that BCG vaccination has the potential to reduce the prevalence of bTB in badgers (Chambers *et al.* 2011, Lesellier *et al.* 2011, Carter *et al.* 2012, Gormley *et al.* 2017). Furthermore although we currently lack empirical estimates of the epidemiological consequences for bTB in cattle, mathematical models predict that badger vaccination, used singly or in combination, should have a beneficial effect (Smith *et al.* 2012, Smith *et al.* 2013, Abdou *et al.* 2016).

Oral delivery of BCG by means of a palatable bait is likely to be the most practical and effective option for mass vaccination of wildlife and an oral TB badger vaccine is currently in development (Chambers 2014). However, there are many challenges to overcome in developing an orally delivered vaccine (Chambers *et al.* 2014) and the requirements for development and authorisation make it likely that a licenced bait will not be available for a number of years. In the meantime, injectable BCG remains the only available option for badger vaccination campaigns.

The present study is the first to estimate vaccine coverage in a badger population subjected to cage trapping and injection. Our analytical approach could also be used to estimate trapping efficiency for other methods of badger bTB control such as culling. We have demonstrated that even at the lower estimate of annual coverage observed, it may be possible to vaccinate in excess of 40% of the badger population in a given year, which is consistent with achieving 70% coverage by the end of a four year annual vaccination campaign.

## Acknowledgements

This study was funded by the Welsh Government. We are indebted to our colleagues in the Welsh Government Office of the Chief Veterinary Officer for their assistance in coordinating the project, and to the OCVO and APHA field staff for carrying out the field work. We are also grateful to the staff of the APHA Central Sequencing Laboratory for expert technical assistance, particularly Saira Crawshaw. Finally, we are indebted to the landowners and tenants in the Intensive Action Area for granting access to their land.

